# Accurate detection and quantification of single-base m6A RNA modification using nanopore signals with multi-view deep learning

**DOI:** 10.1101/2025.08.04.668591

**Authors:** Jun Zhang, Jianbo Qiao, Zhiqun Zhao, Chenglin Yin, Junru Jin, Ding Wang, Wenjia Gao, Leyi Wei

## Abstract

N6-methyladenosine (m6A) is a crucial epitranscriptomic mark. While Nanopore Direct RNA Sequencing (DRS) enables transcriptome-wide detection, most existing methods neglect or lack the capacity to effectively process the intrinsically variable-length raw signals generated by DRS reads.Here, we present MultiNano, a multi-view deep learning framework that converts variable-length raw signals into image-like feature representations, effectively resolving the length inconsistency problem. By integrating raw signal and basecalling features, MultiNano enables accurate and comprehensive transcriptome-wide detection of m6A modifications. Our model achieved state-of-the-art (SOTA) performance in various tasks, including site-level prediction, read-level prediction, cross-species transfer learning, and modification rate estimation. Furthermore, the false positive control strategy implemented in MultiNano significantly enhances the model’s robustness and predictive accuracy, offering a powerful alternative to traditional thresholding-based filtering algorithms. Collectively, our approach provides novel insights for the absolute quantification and single-base resolution of RNA modifications.

## Introduction

First discovered in the 1950s^1^, and to date, RNA modifications are now known to encompass over 150 distinct types^2–4^. Among these, N6-methyladenosine (m6A) stands out as the most abundant and well-characterized internal modification in eukaryotic mRNA^5^. In eukaryotic transcripts, m6A predominantly occurs within the conserved DRACH or RRACH sequence motifs (where D = A, G, or U; R = A or G; H = A, C, or U). This modification is essential for fundamental cellular processes, including mRNA stability^6^, splicing^7^, and translation^8^. For instance, m6A deposition can promote the destabilization of specific mRNAs in embryonic stem cells^9^, and aberrant m6A modification patterns have been linked to a variety of human diseases^10–15^.

In recent years, revolutionary technologies have been developed, enabling a more profound characterization of the complex eukaryotic epitranscriptome^16–21^. Methods such as MeRIP-Seq^22^, m6ACE-Seq^23^, miCLIP^24^, GLORI^25^ relying on antibodies or chemical treatments have been widely employed to profile mRNA modifications. These approaches typically involve inducing changes in modified nucleotides, isolating the RNA, performing reverse transcription, and detecting the resulting signals via short-read cDNA sequencing. While these techniques permit transcriptome-wide mapping of RNA modification sites, they are hindered by inherent limitations, including their reliance on modification-specific antibodies or reagents^26^, and the absence of single-nucleotide resolution^27,28^.

Nanopore Direct RNA sequencing (DRS) technology has opened new avenues for detecting RNA modifications such as m6A^29–31^. In contrast to traditional RNA-seq approaches that necessitate reverse transcription into cDNA, Oxford Nanopore Technologies’ DRS enables the direct sequencing of native RNA molecules, thereby preserving endogenous modifications such as m6A. Each nucleotide, whether modified or not, induces unique alterations in the current signals as RNA molecules pass through the nanopores. These signal perturbations are recorded in real time and used for base identification, with modified bases generating distinguishable deviations from expected patterns. Such alterations can be captured and interpreted using machine learning -based approaches^32,33^, enabling high-resolution detection of RNA modifications^34^.

Broadly, current DRS-based m6A detection approaches fall into two main categories: comparative methods and those based on supervised learning. Comparative methods operate without requiring prior knowledge or training data of known RNA modifications. Instead, they identify statistically significant shifts in signal features by comparing modified samples against control samples presumed to lack modifications. Tools such as Tombo^35^, DRUMMER^36^, Nanocompore^37^, ELIGOS^38^, and xPore^39^ adopt this strategy to infer m6A sites by contrasting the signal profiles of m6A-containing and m6A-depleted samples. Although effective, these methods are significantly limited by their stringent requirement for high-quality m6A-free control samples. Such controls typically necessitate the knock-out (KO) or knock-down (KD) of crucial writer genes like METTL3, which is technically challenging and expensive to produce.

Supervised methods, including EpiNano^40^, nanom6A^41^, m6Anet^42^, and RedNano^43^, employ machine learning to identify m6A sites from Nanopore DRS reads. Typically, these models are trained with labels originating from synthetically modified RNA samples or established experimental protocols, including miCLIP, MeRIP-Seq, and m6ACE-Seq. Among these four representative methods, EpiNano leverages systematic basecalling errors as its primary features. The remaining methods, excluding RedNano, extract features directly from the current signals of Nanopore reads. RedNano, however, uniquely combines features derived from both raw electrical signals and basecalling errors. Despite these varied approaches, supervised methods still face several limitations. A primary limitation is that most existing models do not fully exploit the rich information within indeterminate- length raw electrical signals. They predominantly rely on a restricted set of statistical features, including mean, standard deviation, median, and dwell time. Furthermore, while approaches such as RedNano integrate raw signal data, they often lack a multi-perspective analysis of signal characteristics, which limits their capacity to comprehensively capture the intricate complexity of RNA modifications.

To address these challenges and aim for a highly accurate, biologically applicable approach for Nanopore DRS analysis, we introduce MultiNano. This model integrates basecalling features, raw electrical signals, and signal transformations to construct a multi-view prediction model to, fully leverage the diverse feature representations of Nanopore signal data. By applying MultiNano to human, *Arabidopsis*, and synthetic DRS datasets, we demonstrate its enhanced accuracy in m6A site detection and its direct applicability to biological analyses. Our results show that MultiNano not only outperforms existing approaches in predictive accuracy but also exhibits strong transfer learning capability in cross-species experiments. To the best of our knowledge, MultiNano stands as the first algorithm to introduce a multi-view modeling framework for Nanopore-based m6A detection. Our results clearly indicate that extending this model with additional signal modalities enhances its prediction performance.

## Results

### Boosting m6A site identification from nanopore RNA signals via the MultiNano multi-view learning model

In this study, we developed MultiNano, a multi-view learning model that integrates raw signal and basecalling features. This integration enables a more comprehensive and accurate characterization of m6A modification distribution across multiple species. The MultiNano framework are composed of three main components: the data preprocessing module, the MultiNano core module, and the classification module (Fig.1a). Initially, Nanopore DRS reads are processed to extract relevant features. Basecalling features are fed into a BiLSTM module to capture sequential dependencies, while raw signal features transformed into Gramian Angular Summation Field (GASF) representations^44^ (Fig.1b), and raw signals were processed through a 1D residual networks (ResNet)^45^ module (Fig.1c). These representations are then further analyzed by an optimized ResNet2D module (Fig.1d). This module enhances spatial feature extraction performance by combining channel-wise attention (via SE blocks)^46^ and spatial attention mechanisms^47^. Finally, all features were fused through a fully connected layer. The classification module then employed a multiple instance learning (MIL) strategy^42^ to aggregate read-level methylation probabilities and infered site-level m6A modification probabilities.

**Fig.1.**
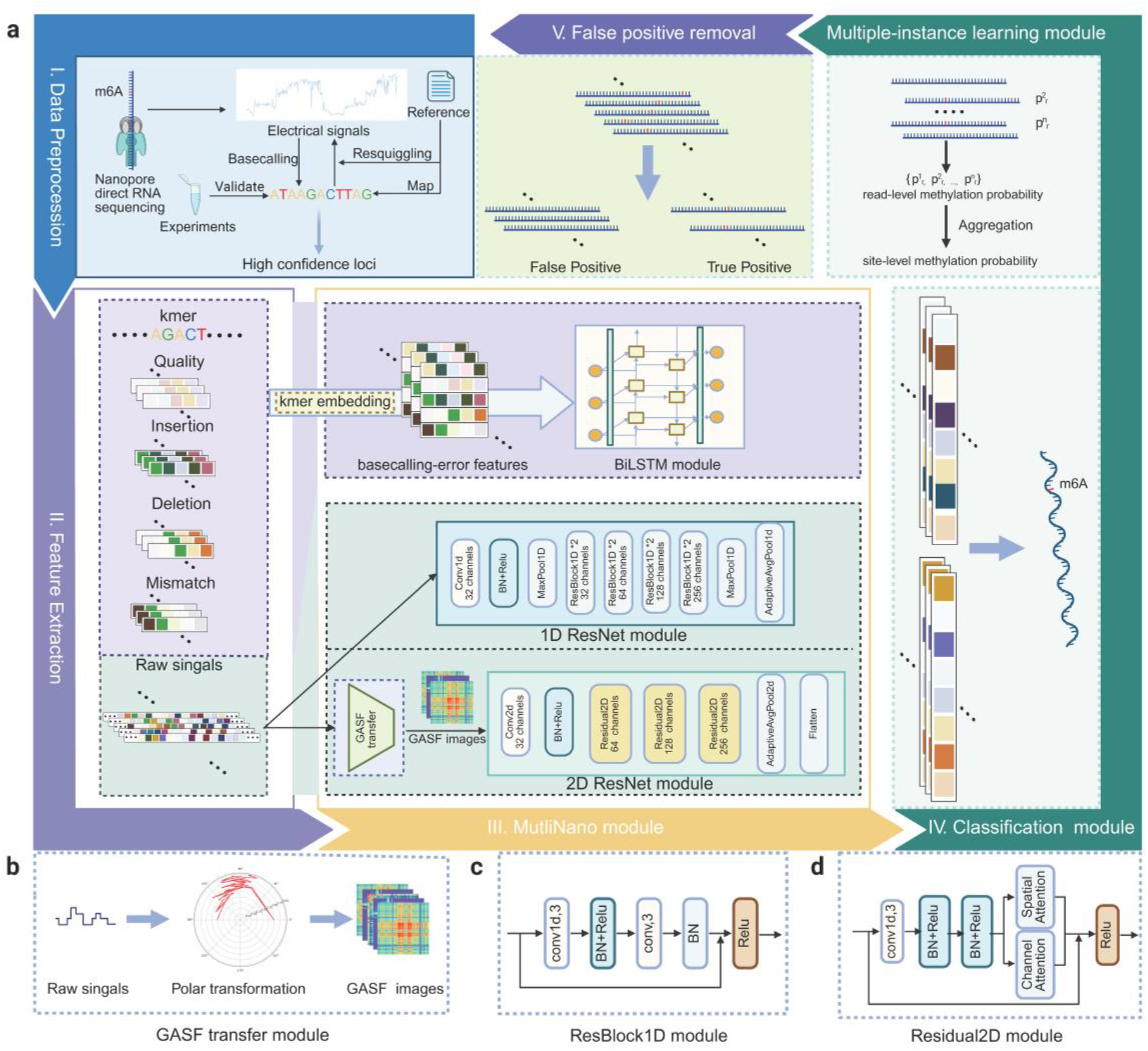
Overview of the MultiNano framework. **a** The overall architecture of the MultiNano framework. **b** Schematic illustration of the GASF transfer module, which is responsible for converting preprocessed signals into image-like representations. **c** Diagram of the ResBlock1D module, the basic building block of the 1D ResNet module, designed to process raw signal information. **d** Diagram of the ResBlock2D module, the basic building block of the 2D ResNet module, designed to process transformed signal representations.

### MultiNano improves site-level methylation prediction

To evaluate MultiNano’s site-level predictive performance for m6A modifications, we tested it on DRS data from the HEK293T dataset^39^. For a fair comparison, all models were retrained from scratch using identical training and validation datasets, and then evaluated in a same testset. The competing supervised methods included EpiNano, nanom6A, m6Anet, and RedNano. Notably, to support all 18 DRACH motifs, we modified the source code of nanom6A which natively supports only 12 RRACH variants without altering its core architecture. MultiNano consistently outperformed other models across all evaluated metrics. On the HEK293T dataset, MultiNano achieved the area under the ROC curve (AUC) of 0.8597 and the area under the precision-recall curve (AUPR) of 0.5448 for DRACH motifs (Fig.2a, Supplementary Table S1). For RRACH motifs, MultiNano reached an AUC of 0.859 and an AUPR of 0.558 (Fig.2b, Supplementary Table S2). These results underscored MultiNano’s robust predictive performance across different motif types. Furthermore, our results indicated that MultiNano outperformed over existing methods across various evaluation criteria, most notably achieving a higher F1 score in identifying m6A sites (Fig.2c,d).To further investigate MultiNano’s ability to predict modification rates under real-world conditions with varying motif proportions, we compared the predicted outputs of all models against the ground-truth motif-level modification distributions. MultiNano achieved the best scores in Pearson correlation (Pearson), mean squared error (MSE), and mean absolute error (MAE). Specifically, its predictions showed the strongest correlation with ground-truth modification proportions, attaining a coefficient of determination (R^2^) of 0.9348, which significantly outperformed RedNano (0.9308), m6anet (0.8616), nanom6a (0.8370), and epinano (0.5364) (Fig.2e, Supplementary Table S3). This confirms MultiNano’s excellent site-level prediction performance and its ability to more accurately reflect the actual modification landscape.

**Fig.2.**
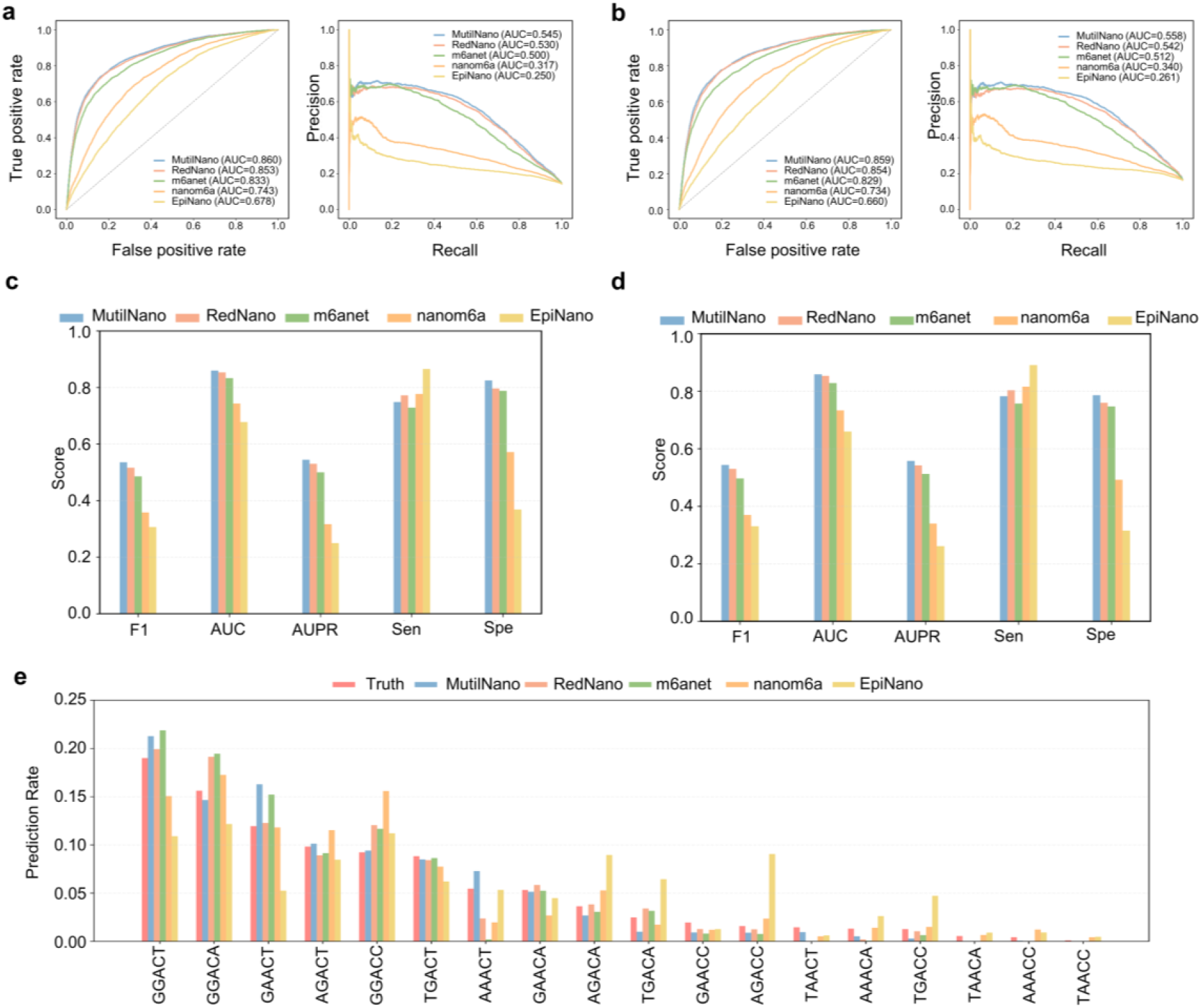
Site-level performance comparison of MultiNano against other models for various motifs in the HEK293T dataset. **a** Performance comparison of MultiNano and other models in detecting m6A sites n the DRACH motif using the HEK293T dataset. **b** Performance comparison of MultiNano and other models in detecting m6A sites on the RRACH motif using the HEK293T dataset. **c** Comparison of the performance metrics of MultiNano and other models on the DRACH motif using the HEK293T dataset. **d** Comparison of the performance metrics of MultiNano and other models on the RRACH motif using the HEK293T dataset. **e** The predicted modification proportions and ground truth across different models on the DRACH motif.

### MultiNano outperforms existing models in read-level methylation prediction

In MultiNano’s classification module, we employed a MIL strategy where read-level methylation predictions serve as the basis for site-level methylation prediction. To evaluate the read-level performance of MultiNano, we benchmarked it against nanom6A and RedNano using both a synthetic RNA dataset^40^ and an in vitro transcription (IVT) dataset^48^. To ensure a fair comparison at the read level, we mirrored our site-level evaluation protocol. All models were trained de novo using identical training and validation sets and subsequently assessed on a unified test set. Notably, to facilitate this direct comparison, we first extended the nanom6A model to support all 18 DRACH motif variants. This comparative analysis revealed MultiNano’s superior performance, achieving an AUC of 0.964 and an AUPR of 0.958 in synthetic RNA dataset (Fig.3a, Supplementary Table S4) and corresponding scores of 0.93 (AUC) and 0.96 (AUPR) in IVT dataset (Fig.3b, Supplementary Table S5), thereby outperforming both nanom6A and RedNano. Importantly, in both datasets, modified bases were consistently predicted with probabilities close to 1, while unmodified bases were predicted with probabilities close to 0 (Supplementary Fig.S1a, b), indicating that most predictions are highly confident and well-calibrated.

**Fig.3.**
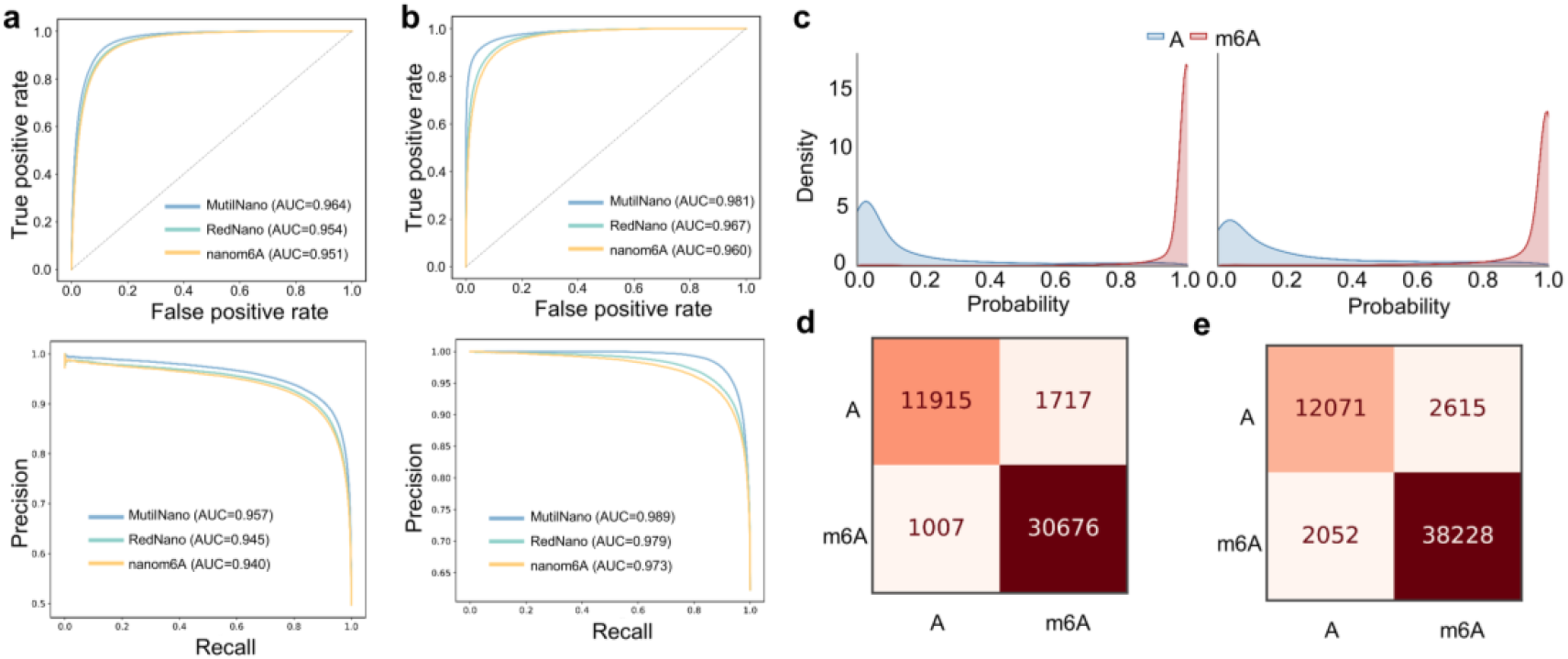
Read-level performance comparison of MultiNano against other models, and MultiNano’s single-site modification identification capability. **a** Comparison of MultiNano and other models at the read-level on the synthetic RNA dataset. **b** Comparison of MultiNano and other models at the read-level on the IVT dataset. **c** Predicted probability distribution for site 1107 (left panel) and site 679 (right panel). **d** Confusion matrix of the prediction results for the site 1107. **e** Confusion matrix of the prediction results for the site 679.

To further assess the model’s ability to discriminate between modified and unmodified states at a single-site resolution, we analyzed two specific loci (base 679 and 1107) on chromosome A2 of the IVT dataset. For both sites, the model assigned probabilities near 1.0 to m6A-modified reads and near 0 to unmodified reads. This high confidence translated into excellent single-site discriminative performance, yielding accuracies of 0.915 and 0.940, and sensitivities of 0.949 and 0.968, respectively (Fig.3d, e). Collectively, these results show MultiNano’s powerful capability to distinguish modification states at the single- site and read level.

### MultiNano demonstrates strong cross-species generalization capability

Supervised learning models fundamentally rely on labeled data for training. However, the scarcity of gold-standard experimental data means that reliable m6A site annotations are unavailable for many species. Therefore, a model’s capacity for cross-species generalization is a crucial concern for its practical utility. To evaluate the cross-species generalizability of the models, we designed a dedicated pipeline. First, MultiNano and its competitors (EpiNano, nanom6A, m6Anet, and RedNano) were independently trained on synthetic RNA and *Arabidopsis* datasets^49^. Subsequently, the performance of these trained models was benchmarked on the human HEK293T dataset to assess their predictive accuracy in a cross-species context. To further test for robustness against real-world data, we also designed a challenging scenario. In this setup, the training sets derived from synthetic RNA and *Arabidopsis* were constructed to have a different distribution of modification motif frequencies compared to the HEK293T train set (Fig.4a). The objective was to assess whether the models could maintain predictive accuracy when trained on varying motif-specific modification stoichiometries. The subsequent evaluation on the HEK293T dataset revealed that MultiNano possessed superior transferability to new species compared to existing methods. When trained on the synthetic RNA datasets, MultiNano demonstrated exceptional performance, achieving an AUC of 0.752 and an AUPR of 0.406. In contrast, the best-performing competing model, RedNano, only reached an AUC of 0.737 and an AUPR of 0.391 (Fig.4b, Supplementary Table S6). This leading performance was further validated on the Arabidopsis datasets, where MultiNano’s AUC and AUPR climbed to 0.833 and 0.508, once again surpassing all competing models, including RedNano (AUC = 0.823, AUPR = 0.505) (Fig.4c, Supplementary Table S7). Taken together, these results demonstrate MultiNano’s ability to accurately identify m6A modification sites, confirming its broad applicability across diverse species. (Fig.4c, Supplementary Table S7). Taken together, these consistent results across diverse datasets emphatically demonstrate MultiNano’s ability to accurately identify m6A modification sites and confirm its broad applicability across various species.

**Fig.4.**
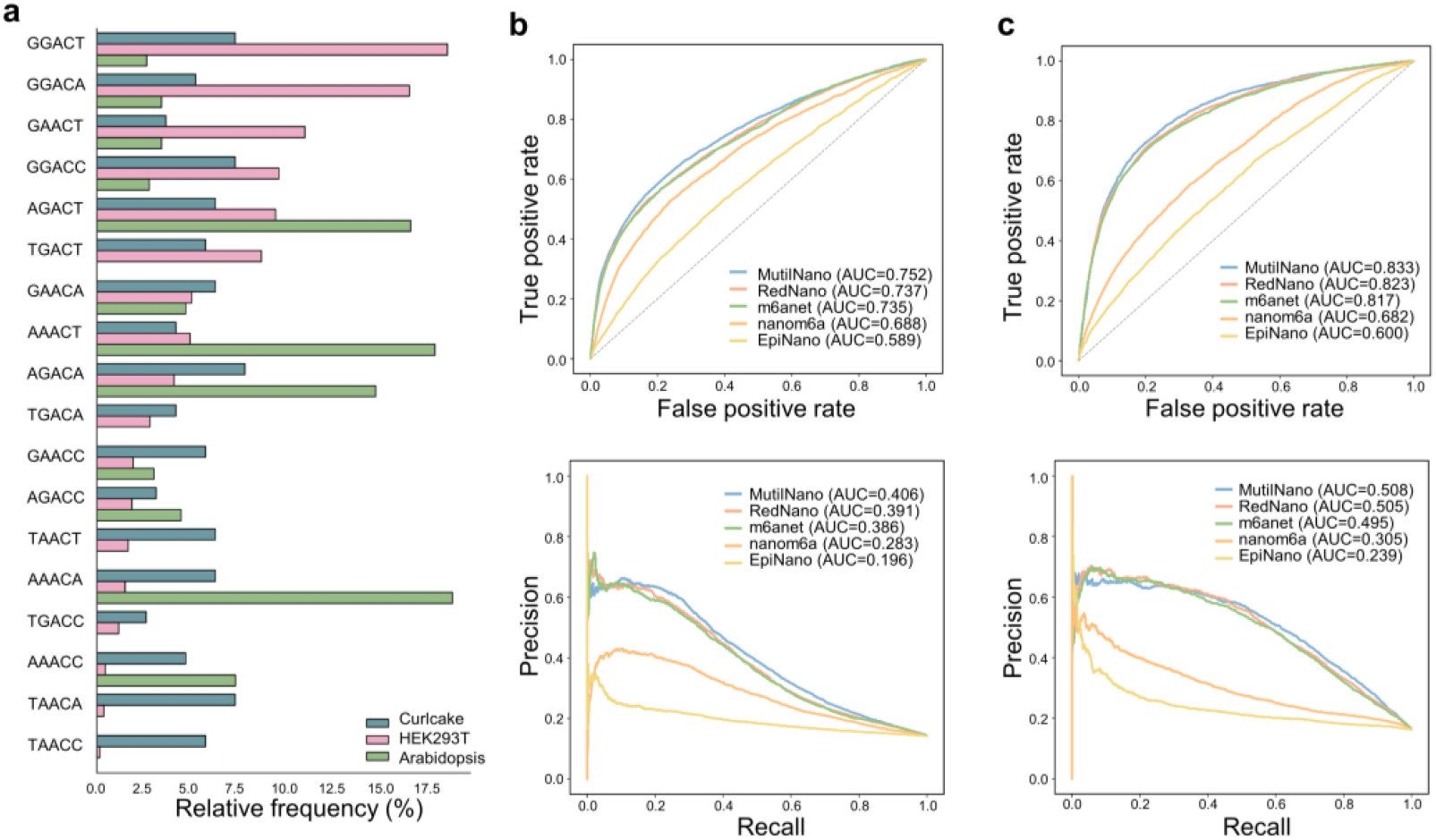
Comparsion of MultiNano and other models in cross-species transfer learning. **a**. Comparisons of the relative frequencies of methylation motifs in the synthetic RNA, *Arabidopsis*, and HEK293T datasets. Each bar indicates the proportion of methylated sites for that motif among all modified 5-mer motifs in the training datasets. **b**. Transfer learning performance from the synthetic RNA dataset to the HEK293T dataset, measured by AUC and AUPR. **c**. Transfer learning performance from the *Arabidopsis* dataset to the HEK293T dataset, measured by AUC and AUPR.

### MultiNano enables precise estimation of RNA modification rates

Modification of m6A is a dynamic and reversible RNA modification with diverse functions in gene expression regulation^50^. Quantifying m6A can reveal its dynamic changes, for instance, under stress conditions like hypoxia^51^ and heat shock^52^. Despite the development of various m6A mapping tools, current methods face limitations in achieving absolute quantification and single-base resolution^25^. To address this, we designed experiments where benchmark DRS datasets were generated with precisely controlled modification rates, by mixing modified and unmodified reads from synthetic RNA and IVT sources across the full stoichiometric range (0% to 100%). On this basis, we evaluated MultiNano’s ability to quantify varying modification stoichiometries.

Firstly, predictions on these two datasets showed a high correlation between our predictions and the ground truth: an R^2^ of 0.886 and a Pearson correlation of 0.951 on the synthetic RNA dataset, and an R^2^ of 0.907 and a Pearson correlation of 0.957 on the IVT dataset. Furthermore, the results were highly enriched near the ground truth, with a sharp decrease in quantity further away and zero instances at the furthest points, demonstrating the reliability of our model’s predictions (Fig.5a, b). To further assess the deviation between our model’s predictions and the ground truth, we calculated the differences. The results indicated that the differences between our model’s predictions and the ground truth, across the full stoichiometric range (0% to 100%), were highly concentrated around zero (Fig.5c, d). This further validated the robustness and accuracy of the model in predicting modification ratios. Moreover, compared to existing models like RedNano (Supplementary Fig.3a-f) and nanom6A (Supplementary Fig.4a-f), our model outperformed them across all metrics in both datasets (Supplementary Table. S8,9), offering new insights and crucial model support for modification rate prediction.

**Fig.5.**
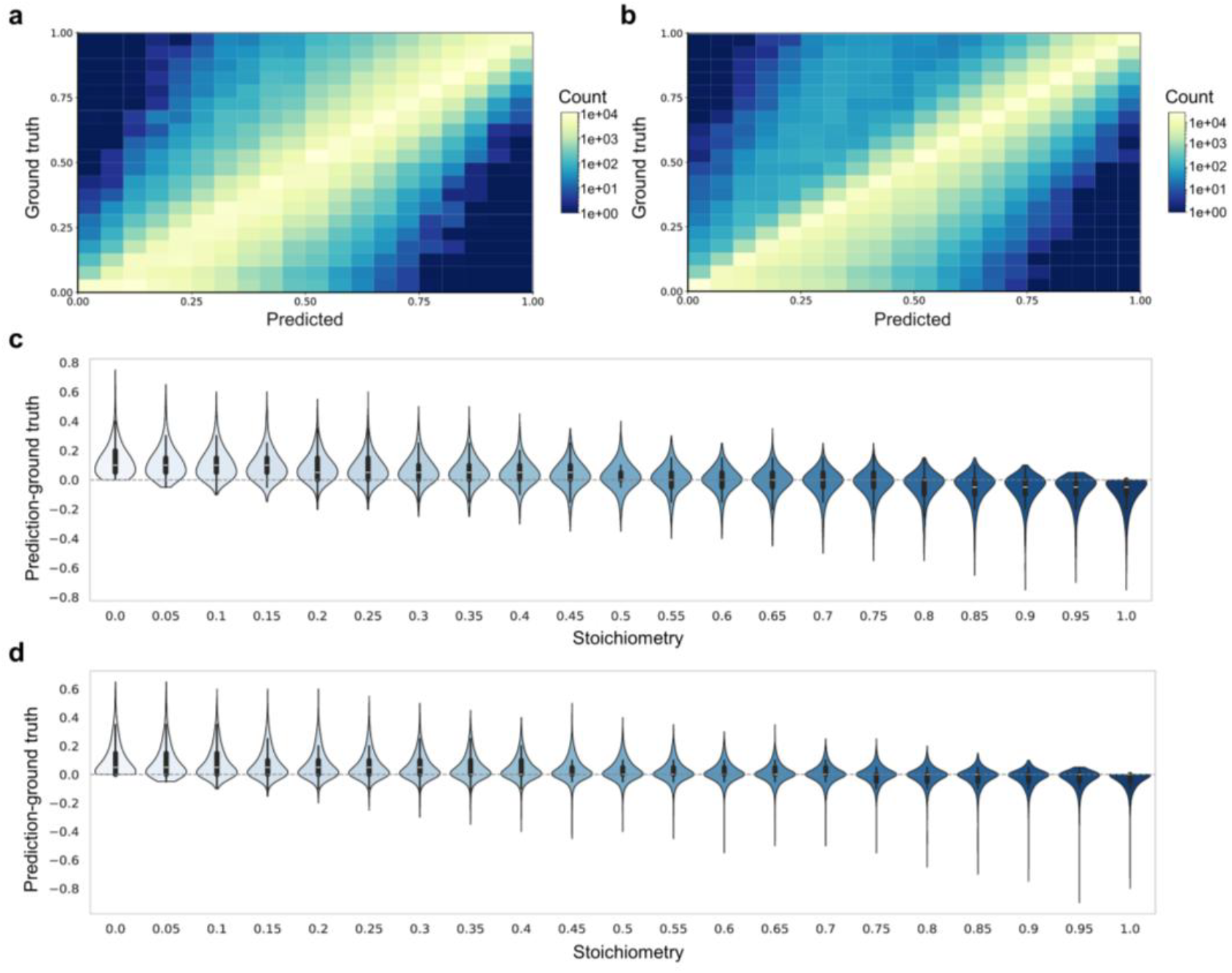
MultiNano’s capability for estimating modification rates. **a** Correlation between predicted methylation stoichiometry by MultiNano and the ground truth in the synthetic RNA dataset. **b** Correlation between predicted methylation stoichiometry by MultiNano and the ground truth in the IVT dataset. **c** Deviation between predicted and ground truth methylation stoichiometry in the synthetic RNA dataset. **d** Deviation between predicted and ground truth methylation stoichiometry in the IVT dataset.

### MultiNano effectively suppresses false positive predictions

A major limitation of current supervised learning models for methylation prediction is their high rate of false positives, which severely impairs predictive reliability. To counteract this, a common practice is to apply a high-confidence threshold (e.g., a cutoff of 0.9)^53^ to filter predictions. While this method effectively reduces false positives, it introduces a critical trade-off, as a vast number of true positive sites with prediction scores below the cutoff are inadvertently discarded, leading to a substantial loss in recall.

To address this critical issue, we built our approach on the key observation that true and false positive sites display distinguishable patterns in their underlying read-level probability distributions (Fig.6a, b). To harness these patterns, we engineered a comprehensive set of 29 statistical features for each site based on its collection of read-level probabilities. (Supplementary Data1, Supplementary Fig.S5, 6) This feature set was designed to capture properties including central tendency and dispersion (e.g., minimum and maximum values, standard deviation), quantiles, shape characteristics (e.g., skewness, kurtosis), and sequential trends. To evaluate the differences between the two groups, we note that the large sample size may lead to statistically significant p-values that lack practical significance. Therefore, we introduced Cliff’s delta^54^ (Cliff’s d) as a non-parametric effect size measure to quantitatively assess the magnitude of the difference between the false positive and true positive. Finally, we developed a novel XGBoost-based algorithm^55^ trained on these features to accurately filter out false positive predictions.

**Fig.6.**
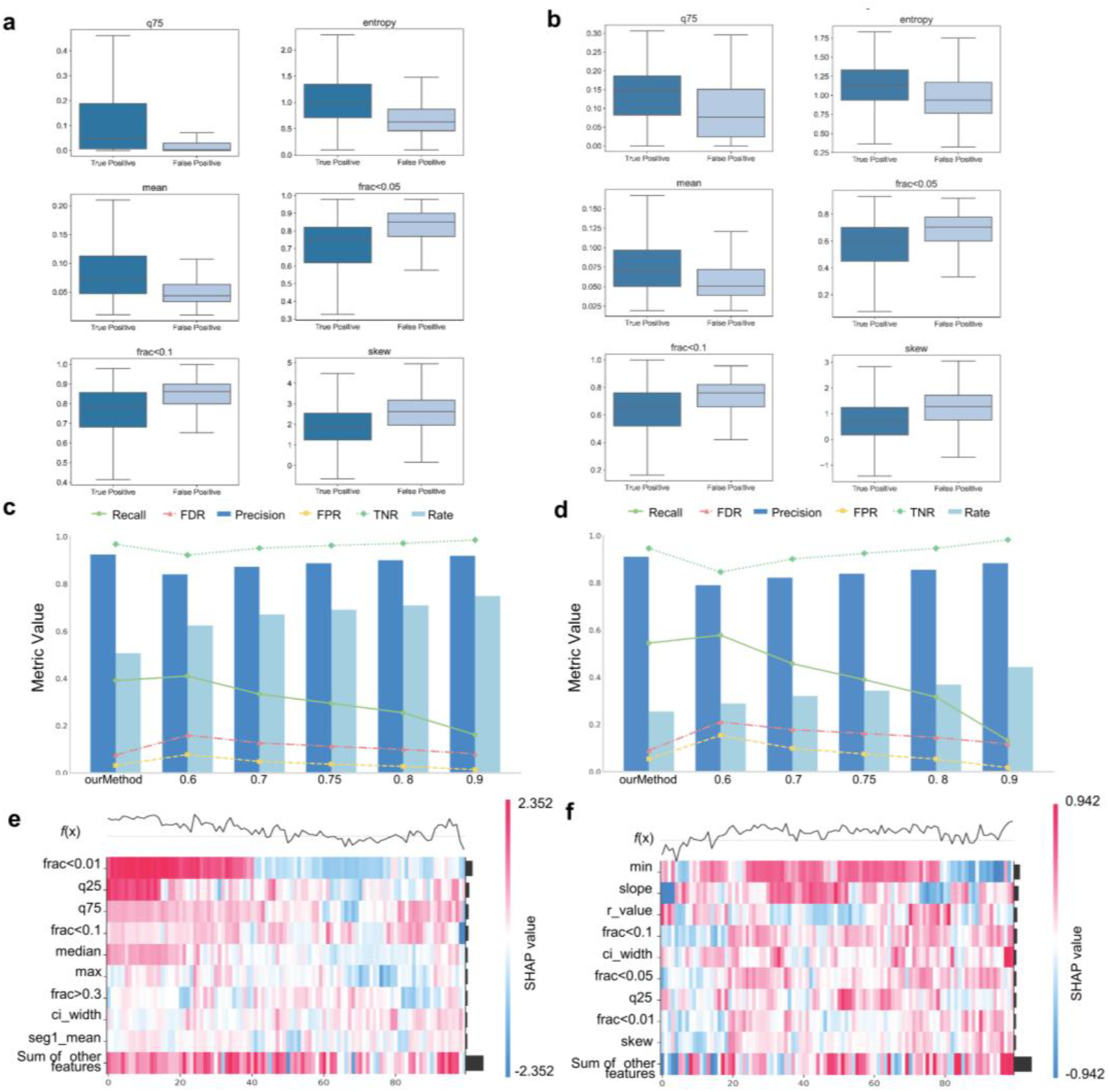
Implementation of site-level false positive control via read-level modification probability distributions. **a, b** Comparison of predicted read-level methylation distributions between true positives and false positives when the model is trained on the synthetic RNA dataset (a) or the *Arabidopsis* dataset (b) and tested on the HEK293T dataset. **c, d** Performance comparison between the conventional thresholding-based approach and our false positive control module when transferring from the synthetic RNA dataset (c) or the *Arabidopsis* dataset (d) to the HEK293T dataset. **e, f** SHAP heatmaps illustrating model interpretability under false positive control, trained on the synthetic RNA dataset (e) and the *Arabidopsis* dataset (f), respectively.

The robustness of this approach was evaluated in two scenarios: applying an *Arabidopsis*- trained model to a human HEK293T dataset, and applying a synthetic RNA-trained model on the same human dataset. In both independent tests, our method demonstrated its ability to resolve the classic precision-recall trade-off. It achieved an exceptional precision of over the method based thresholding with cutoff of 0.9, registering scores of 0.910 and 0.924 (Supplementary Table S10, 11), respectively, while keeping a recall much higher than that achieved by simpler high-thresholding with cutoff 0.7 (Fig.6c, d). This superior balance ultimately provides a more effective means to enhance model reliability, accurately purging false positives while minimizing the loss of true modification signals.

To further interpret the decision-making process of our XGBoost model and validate the utility of our engineered feature set, we employed SHapley Additive exPlanations (SHAP)^56^. The analysis revealed the relative importance of 29 features in discriminating between true and false positive sites (Supplementary Fig.S7a, b). As anticipated, this analysis identified the mean, standard deviation, and range of read-level probabilities as the most influential features. Notably, the analysis also confirmed that a variety of features, including those related to distribution shape and sequential trends, made meaningful, non-redundant contributions to the final prediction (Fig.6e, f). In conclusion, our comprehensive feature engineering strategy proved to be sound, as it effectively leverages a combination of complementary feature types to achieve its high predictive accuracy.

### Ablation study demonstrates the effectiveness of each component within MultiNano

MultiNano’s architecture synergistically integrates three distinct feature modalities: basecalling, raw electrical signals, and transformed signal characteristics. To systematically evaluate the contribution of each modality, we conducted a comprehensive ablation study. Site-level evaluations were performed on both the *Arabidopsis* and synthetic RNA datasets, comparing models trained with single, dual, and all three feature types.

The results revealed a clear performance hierarchy. On the *Arabidopsis* dataset, the full model incorporating all three feature types achieved an AUC of 0.921, representing a performance gain of at least 2% over any dual-feature combination. This synergistic effect was consistently observed on the synthetic RNA dataset. Furthermore, across both datasets, all dual-feature models significantly outperformed their single-feature counterparts. These findings strongly indicate that each feature modality captures unique and complementary information essential for robust m6A prediction (Table1).

**Table 1.** Blation study of model performance using three types of features: basecalling features (basecalling), raw signals (raw), and transformed signals (signal), as well as their combinations.

| Data | Combine | Feature | F1 | AUC | AUPR | Sensitivity | Specificity |
| --- | --- | --- | --- | --- | --- | --- | --- |
| *Arabidopsis* | Single | basecalling | 0.800 | 0.876 | 0.871 | 0.801 | 0.797 |
|  |  | raw | 0.791 | 0.875 | 0.881 | 0.873 | 0.664 |
|  |  | signal | 0.719 | 0.807 | 0.814 | 0.705 | 0.746 |
|  | Dual | basecalling_raw | 0.818 | 0.903 | 0.913 | 0.818 | 0.818 |
|  |  | basecalling_signal | 0.799 | 0.879 | 0.887 | 0.825 | 0.760 |
|  |  | raw_signal | 0.830 | 0.898 | 0.914 | 0.845 | 0.808 |
|  | Triple | all | 0.857 | 0.919 | 0.929 | 0.876 | 0.837 |
| Curlcake | Single | basecalling | 0.976 | 0.994 | 0.995 | 0.974 | 0.979 |
|  |  | raw | 0.906 | 0.969 | 0.971 | 0.958 | 0.843 |
|  |  | signal | 0.891 | 0.948 | 0.947 | 0.895 | 0.885 |
|  | Dual | basecalling_raw | 0.968 | 0.991 | 0.993 | 0.958 | 0.979 |
|  |  | basecalling_signal | 0.957 | 0.995 | 0.995 | 0.937 | 0.979 |
|  |  | raw_signal | 0.933 | 0.982 | 0.983 | 0.953 | 0.911 |
|  | Triple | all | 0.976 | 0.994 | 0.995 | 0.974 | 0.979 |

Subsequently, we performed a comparative analysis to identify the optimal signal-to-image transformation strategy within MultiNano’s image conversion module. We evaluated three established techniques for encoding time-series data as images: the GASF, the Gramian Angular Difference Field (GADF), and the Recurrence Plot (RP). We benchmarked models using eatch techniques individually and in combination on both the *Arabidopsis* and synthetic RNA datasets. Results showed that the the GASF-based model consistently outperformed models all other configurations. This finding led us to select GASF as the definitive image transformation strategy for MultiNano (Table2).

**Table 2.** Ablation study of model performance using three image transformation methods:GASF, GADF, and RP, as well as their combinations.

| Dataset | Combine | Transform | F1 | AUC | AUPR | Sensitivity | Specificity |
| --- | --- | --- | --- | --- | --- | --- | --- |
| *Arabidopsis* | Single | GASF | 0.857 | 0.919 | 0.929 | 0.876 | 0.832 |
|  |  | RP | 0.821 | 0.912 | 0.919 | 0.904 | 0.701 |
|  |  | GADF | 0.837 | 0.921 | 0.930 | 0.863 | 0.801 |
|  | Dual | GASF_RP | 0.845 | 0.909 | 0.911 | 0.859 | 0.825 |
|  |  | GADF_GASF | 0.787 | 0.855 | 0.861 | 0.839 | 0.708 |
|  |  | GADF_RP | 0.824 | 0.912 | 0.919 | 0.811 | 0.842 |
|  | Triple | GADF_GASF_RP | 0.833 | 0.900 | 0.911 | 0.863 | 0.790 |
| Curlcake | Single | GASF | 0.976 | 0.994 | 0.995 | 0.974 | 0.979 |
|  |  | RP | 0.920 | 0.994 | 0.994 | 0.990 | 0.838 |
|  |  | GADF | 0.964 | 0.994 | 0.994 | 0.979 | 0.948 |
|  | Dual | GASF_RP | 0.956 | 0.987 | 0.987 | 0.963 | 0.948 |
|  |  | GADF_GASF | 0.932 | 0.989 | 0.989 | 0.963 | 0.895 |
|  |  | GADF_RP | 0.919 | 0.987 | 0.988 | 0.979 | 0.848 |
|  | Triple | GADF_GASF_RP | 0.959 | 0.989 | 0.989 | 0.979 | 0.937 |

## Discussion

Current methods for Nanopore-based m6A detection struggle to fully utilize the rich, variable-length raw electrical signals, often relying on limited statistical or basecalling error features^30,40–42,48,57,58^. To overcome this, we developed MultiNano, a multi-view deep learning framework with a central innovation:the transformation of 1D raw signals into 2D image-like representations via GASF. This novel approach not only resolves the issue of inconsistent signal length but also enables powerful neural networks to capture complex temporal patterns. By synergistically integrating these image-based features with raw signal features and basecalling error profiles, MultiNano constructs a more holistic and robust representation of m6A modification events.

Our comprehensive ablation studies confirmed the efficacy of this multi-view design, revealing a clear performance hierarchy where the GASF configuration significantly surpassed all other tested setups, including single, dual, or hybrid combinations of image transformations. This provides strong evidence that each feature modality captures unique information; the raw signal reflects physical perturbation, basecalling represent downstream algorithmic impact, and GASF-encoded images capture higher-order temporal patterns. This result validates GASF as our optimal choice for the image conversion module, and this synergistic integration is the primary reason for MultiNano’s leading performance in both site-level and read-level predictions across multiple datasets.A key contribution of our work is the development of a novel, feature-based false positive filtering strategy that directly confronts the limitations of conventional high-threshold methods. While high-threshold cutoffs invariably force a trade-off between precision and recall^53^, our XGBoost-based module successfully resolves this dilemma. Our solution is learning the subtle distributional differences between true and false positive sites. And our results demonstrate that MultiNano can achieve high precision (>0.9) while simultaneously maintaining a high recall rate, thereby significantly enhancing model reliability. Furthermore, MultiNano’s strong performance in cross-species experiments underscores its high generalizability, making it a practical tool for m6A study in non-model organisms where high-quality training data is scarce.

Despite its robust performance, MultiNano has several limitations that warrant consideration. First, the multi-view architecture, particularly the image conversion and ResNet2D components, is more computationally intensive than methods relying solely on statistical features. Second, like all supervised methods, MultiNano’s performance is inherently dependent on the quality of the upstream data processing, including the accuracy of basecalling (Guppy) and signal-to-reference alignment (Tombo re-squiggle). This also means that future improvements in these foundational tools are expected to further enhance MultiNano’s predictive power. Finally, while the framework is general, applying it to other types of RNA modifications would necessitate the generation of new, high-quality labeled datasets for model training.

Looking ahead, several exciting avenues exist for future work. A crucial next step is to adapt and validate MultiNano on data generated using newer Oxford Nanopore Technologies kits, such as the recent RNA004 release. While this kit significantly improves basecalling accuracy, it may also alter the raw signal distributions, necessitating model re- training or fine-tuning to take full advantage of this new technology. A more ambitious goal is to extend the framework to distinguish between different modification types on the same base (e.g., telling apart unmodifined, m1A and m6A on an adenosine). This would elevate MultiNano from a binary detector to a true multi-class classifier, offering unprecedented resolution.Sucess here will require the generation of high-quality training datasets with simultaneous labels for each modification type. Finally, model’s core multi-view architecture can be optimized by exploringmore efficient deep learning structures, further broadening its application for large-scale biological investigations.

In conclusion, MultiNano establishes a new benchmark for m6A detection from Nanopore DRS. By synergistic feature integration whthin a novel multi-view framework and an advanced false-positive filtering strategy, MultiNano provides superior accuracy, robustness, and broad application for epitranscriptomic research.

## Methods

### Processing and feature extraction of DRS data DRS data and reference

All DRS datasets utilized in this study were obtained from previously published references. For model training, we downloaded two replicates (rep1 and rep2) of the HEK293T cell line. An independent replicate (rep3) from the same cell line was reserved for model evaluation. For the human datasets, we used GRCh38 Ensembl release version 91 as the reference genome and transcriptome. Regarding the synthetic curlcake datasets, replicate 1 (rep1) was used for training, while replicate 2 (rep2) was used for evaluation. The corresponding reference sequences of the synthetic RNA were obtained from reference^40^. For the *Arabidopsis* datasets, replicates 1 and 2 were used for training, and replicate 3 was used for evaluation. The TAIR10 reference genome and Araport11 gene annotation were retrieved from National Center for Biotechnology Information (NCBI)^59^ and Araport11^60^, respectively.

### Basecalling, re-squiggling, and read mapping

Prior to feature extraction, we performed basecalling, re-squiggling, and mapping on all DRS datasets. For the human, curlcake, and *Arabidopsis* datasets, basecalling was conducted using Guppy (v3.1.5)^61^. Second, We then mapped the raw current signals of each read to the corresponding contiguous bases in the reference transcriptome using the re-squiggle module from Tombo (v1.5.1)^35^. Subsequently, minimap2 (v2.17)^62^ was employed to align each read to the reference transcriptome. Downstream processing of Sequence Alignment/Map (SAM/BAM) files, including sorting and indexing, was performed using Samtools (v1.7)^63^.

### Feature extraction

We searched for the target base A within DRACH/RRACH motifs and extracted both signal-based and basecalling-error features at each candidate site. For signal features, we followed the same preprocessing strategy as RedNano^43^. Given the variable length of raw current signals, we set a fixed threshold of 65 time points. Signals shorter than 65 were padded with zeros at both ends, while signals longer than 65 were randomly downsampled to 65 points. For basecalling-error features, we first converted the read-to-reference alignments into tab-delimited format using the sam2tsv module from JVarkit (https://github.com/lindenb/jvarkit). For each 5-mer centered at the target base, we extracted the alignment status (match, mismatch, insertion, and deletion) and the base quality score of each nucleotide in the 5-mer.

### GASF transformation

After preprocessing the raw current signals, we apply the GASF transformation in the signal transformation module. GASF maps a time series into the polar coordinate space and constructs a symmetric image matrix by computing the cosine of the sum of angles. Given a signal sequence

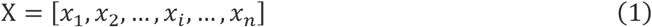

where *x*_i_denotes the current signal at position i, we first remove the padding-0 introduced during preprocessing. The sequence is then normalized as follows:

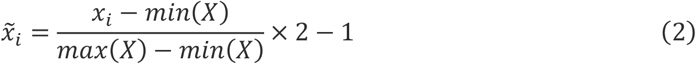

where 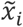represents the normalized value of signal *x*_i_, *min*(*??*) and *m??x*(*??*) denote the minimum and maximum values of the signal sequence.

Then, we perform polar coordinate mapping:

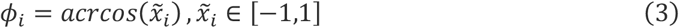

where 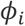 is the angle corresponding to the normalized signal 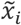. Next, we compute the GASF image matrix using:

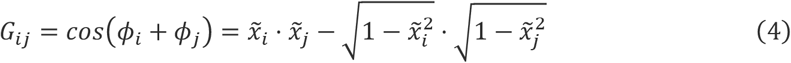

where each element *G*_*ij*_ captures the cosine of the sum of angles between signal positions iand j, quantifying their pairwise relationship:

Finally, we obtain the GASF matrix G as:

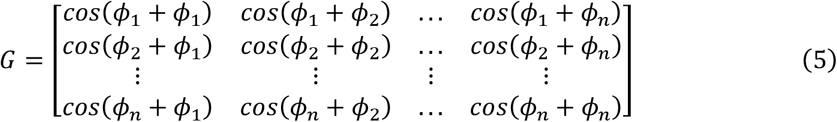

This transformation provides a 2D representation of signal sequences, capturing temporal dynamics through angular summation relationships.

### Attention mechanisms in GASF feature enhancement

To better exploit the visual characteristics of GASF representations, we enhanced the ResNet2D module by incorporating both spatial and channel attention mechanisms.

### Spatial attention

Given an input GASF feature map ?? ∈ ℝ^*C*×*H*×*W*^, we first perform average pooling and max pooling along the channel dimension to generate two spatial descriptors:

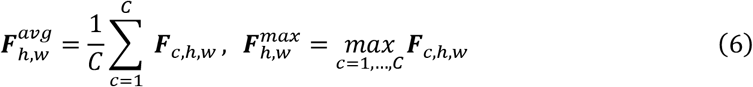

These two descriptors, 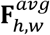 and 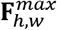, capture complementary spatial information. We concatenate them along the channel axis and apply a convolutional operation with a 7 × 7 kernel followed by a sigmoid activation to obtain the spatial attention map:

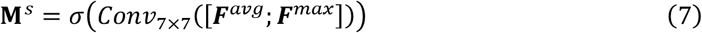

This spatial attention map **M**^s^ ∈ ℝ^1×H×W^highlights the importance of each spatial location. The final spatially attended feature map is obtained via element-wise multiplication:

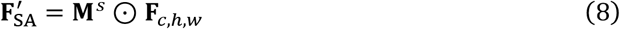

### Channel attention

To further enhance the representational capacity across channels, we apply a channel attention mechanism. We begin by computing global average pooling and global max pooling along the spatial dimensions for each channel:

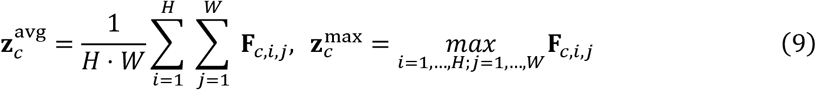

These descriptors, **z**^avg^, **z**^max^ ∈ ℝ^C^, summarize the channel-wise information. We concatenate them and pass the result through a shared two-layer fully connected (FC) network to generate channel attention weights:

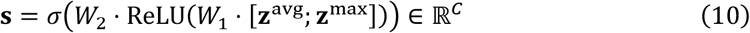

The resulting channel attention weights **s** indicate the importance of each channel. These weights are broadcast across the spatial dimensions and applied to the input feature map via channel-wise multiplication:

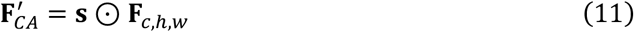

### Combined attention

Finally, we combine both spatial and channel attention outputs to form a joint attention-enhanced representation:

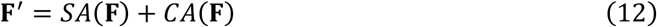

where 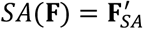 and 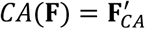. The resulting feature map **F**^′^ integrates both spatial and channel attention, allowing the network to focus on the most informative regions and channels within the GASF representation.

### Multiple-instance learning module

In this study, we formulate the detection problem as a MIL task, a strategy inspired by m6Anet. Each candidate site is treated as a “bag,” comprising multiple “instances” (individual reads). While site-level (bag-level) labels are available (indicating whether a site is modified or not), the read-level (instance-level) labels remain unknown. This weakly supervised setting is particularly well-suited for nanopore signal data, as only a subset of reads may exhibit modification signals.

### Instance encoding

Given a bag *B* = {*x*_1_, *x*_2_, …, *x*_k_}, each instance *x*_j_ is processed by a shared feature extractor 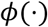, which is implemented using a neural network that integrates Basecalling features, raw signal features, and their corresponding GASF representations. This yields instance-level embeddings:

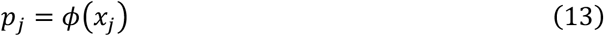

### Instance aggregation

To obtain a bag-level representation, we aggregate the embeddings {z_1_, z_2_, …, z_k_} using a permutation-invariant operator. We apple a standard Noisy-OR aggregation defined as:

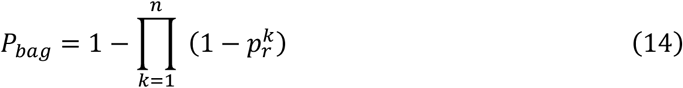

where *p*_j_ is the instance-level probability, *P*_*bag*_ is the bag-level probability.

To simplify the MIL problem, we adopt the cross-entropy function as our training loss.

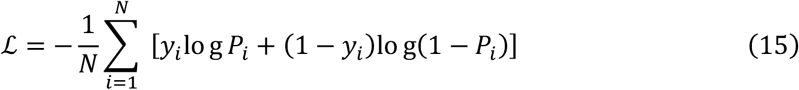

### Model training and evaluation

### Ground truth labels of modified sites

For the curlcake dataset, we utilized all RRACH/DRACH motif sites from both unmodified and m6A-modified Nanopore reads as the benchmark set of candidate sites. The same approach was applied to IVT dataset. For the *Arabidopsis* dataset, we considered sites detected in VIRc wild-type DRS reads as modified, and those detected in the vir-1 mutant as unmodified^49^. For the human dataset, we used m6A-modified sites identified in HEK293T cells based on both m6ACE-seq^39^ and miCLIP data^24^. We included METTL3- dependent sites that exhibited a methylation fold-change (WT/KO) ≥ 4.0 and a one-sided t-test p-value < 0.05 in the m6ACE-seq dataset.

### Model performance evaluation metrics

To evaluate the performance of our model, we utilized five standard metrics: F1 score, AUC, AUPR, Sensitivity (Sen), and Specificity (Spe)^64–66^.

The ROC curve is plotted by comparing the True Positive Rate (TPR) and False Positive Rate (FPR), defined as:

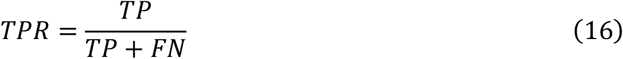

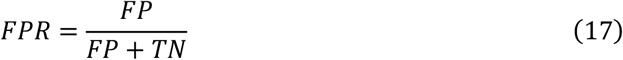

The PR curve is constructed using Precision and Recall, where:

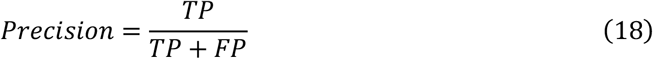

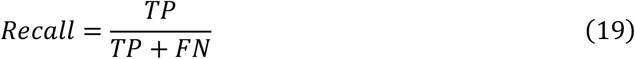

The other evaluation metrics are defined as follows:

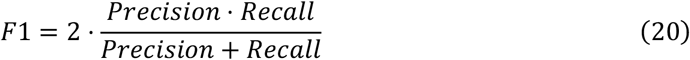

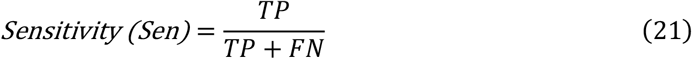

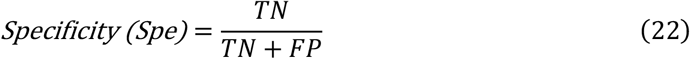

To evaluate the goodness of fit between our model’s predictions and the ground truth, we adopted four metrics: the Pearson, MAE, MSE, and R^2^. These four metrics are defined as follows:

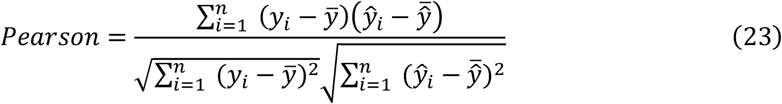

Where ŷ_i_represents the model’s predicted value for the i-th sample, *y*_i_represents the ground truth value for the i-th sample, 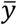 represents the mean of all ground truth values, n represents the total number of samples. These definitions also apply to the following metrics:

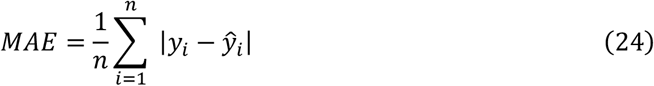

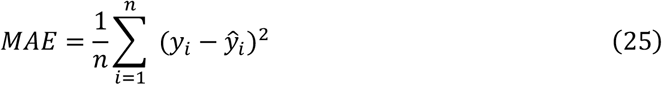

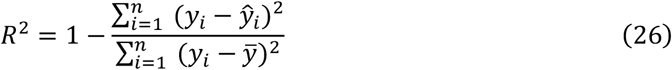

And the definition of Cliff’s d is as follows:

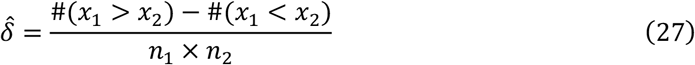

Where #(*x*_1_ > *x*_2_) represents the total number of pairs where a value from group 1 is greater than a value from group 2, #(*x*_1_ < *x*_2_) represents the total number of pairs where a value from group 1 is less than a value from group 2, *n*_1_ is the sample size of group 1, *n*_2_ is the sample size of group 2.

And the Cliff’s Delta effect size (*|δ*|) was interpreted as negligible if |*δ*|<0.147, small if 0.147≤|*δ*|<0.33, medium if 0.33≤|*δ*|<0.474, and large if |*δ*|≥0.474^67^.

### Comparison with other models for m6A detection

#### EpiNano

We employed EpiNano v1.2 (available at https://github.com/novoalab/EpiNano) to extract basecalling-derived features for m6A detection. Specifically, we collected the mean base quality, mismatch frequency, insertion frequency, and deletion frequency for each base within the DRACH motif. Additionally, we utilized the Support Vector Machine (SVM) models provided by EpiNano to perform site-level classification of m6A modifications.

#### Nanom6A

We utilized nanom6A (available at https://github.com/gaoyubang/nanom6A) for m6A site prediction. The model employs an XGBoost classifier trained on features derived from normalized raw signals, including the median, mean, standard deviation, and dwell time of current intensities.

#### m6Anet

We employed m6Anet (available at https://github.com/GoekeLab/m6anet) for m6A site prediction. The Nanopore DRS reads were preprocessed using Nanopolish (v0.14.1)^68^, following the official m6Anet pipeline.

#### RedNano

We utilized RedNano (available at https://github.com/Derryxu/RedNano) for m6A site prediction, following the official usage guidelines. The model integrates raw signal features and basecalling-error features using a deep learning framework tailored for DRS data.

## Data availability

All sequencing data generated in this study are publicly available. The DRS reads of synthesized RNA and the in vitro transcription dataset have been deposited in the National Center for Biotechnology Information (NCBI) under accessions PRJNA511582 and SRP166020, respectively. The DRS reads of Arabidopsis vir-1/VIRc lines and human wild-type HEK293T cells are available in the European Nucleotide Archive (ENA) under accessions PRJEB32782 and PRJEB40872, respectively.

## Code availability

The source code of the MultiNano is freely available for research purposes at Github: https://github.com/zhangjun640/MultiNano.

## Acknowledgements

The work was supported by the Natural Science Foundation of China (No. 62322112) and the Science and Technology Development Fund of Macao (No. 0133/2024/RIB2).

