## Supplementary information for "Accurate detection and quantification of single-base m6A RNA modification using nanopore signals with multi-view deep learning"

### Supplementary Figures

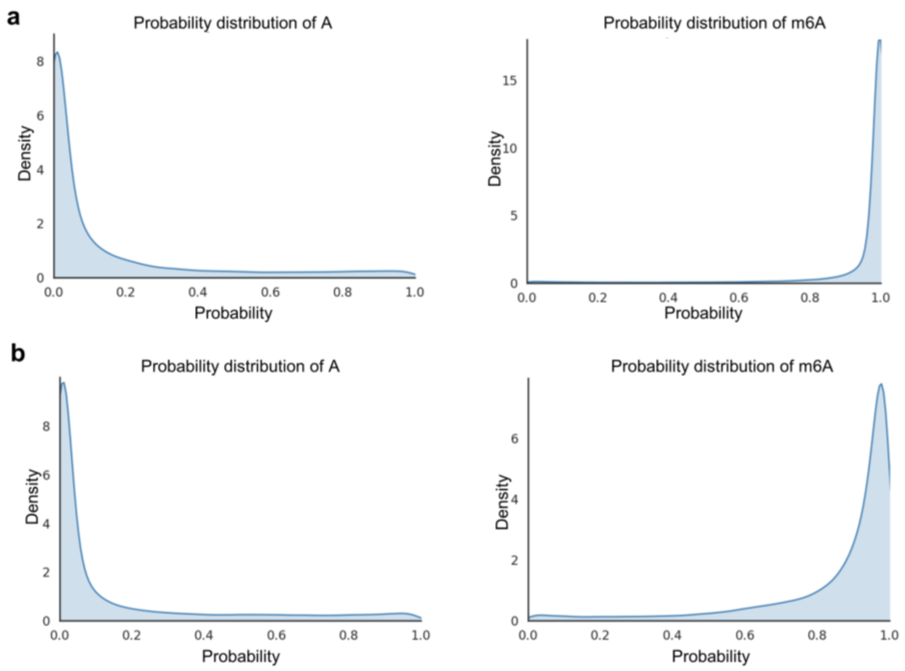


**Supplementary Fig.1 Distribution of predicted methylation probabilities. a,b** Distribution of predicted methylation probabilities for modified and unmodified sites. Predicted probability distributions are shown for models trained on the synthetic RNA dataset (a) and the IVT dataset (b), comparing modified versus unmodified sites.

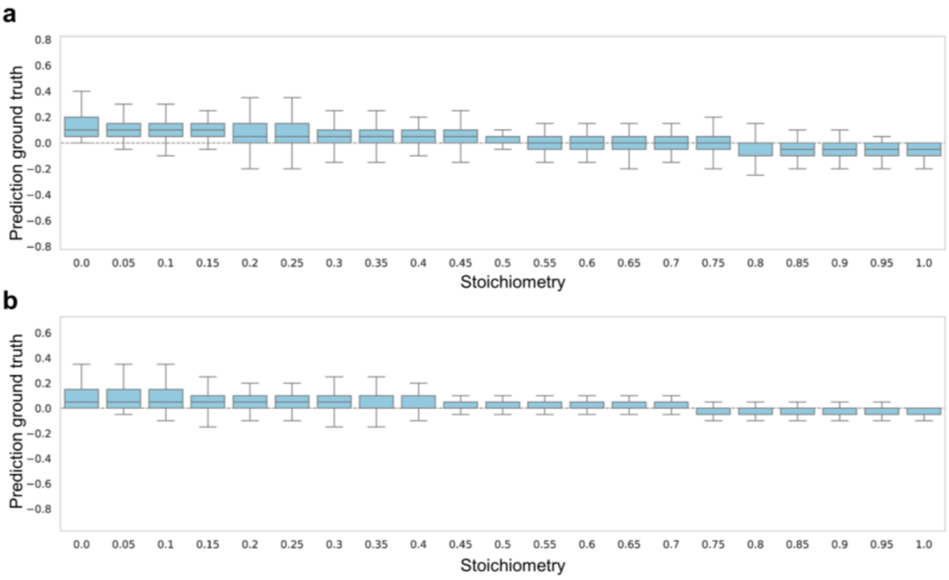


**Supplementary Fig.2 The difference between predicted and ground truth. a** Deviation between predicted and ground truth methylation stoichiometry in the synthetic RNA dataset. **b** Deviation between predicted and ground truth methylation stoichiometry in the IVT dataset.

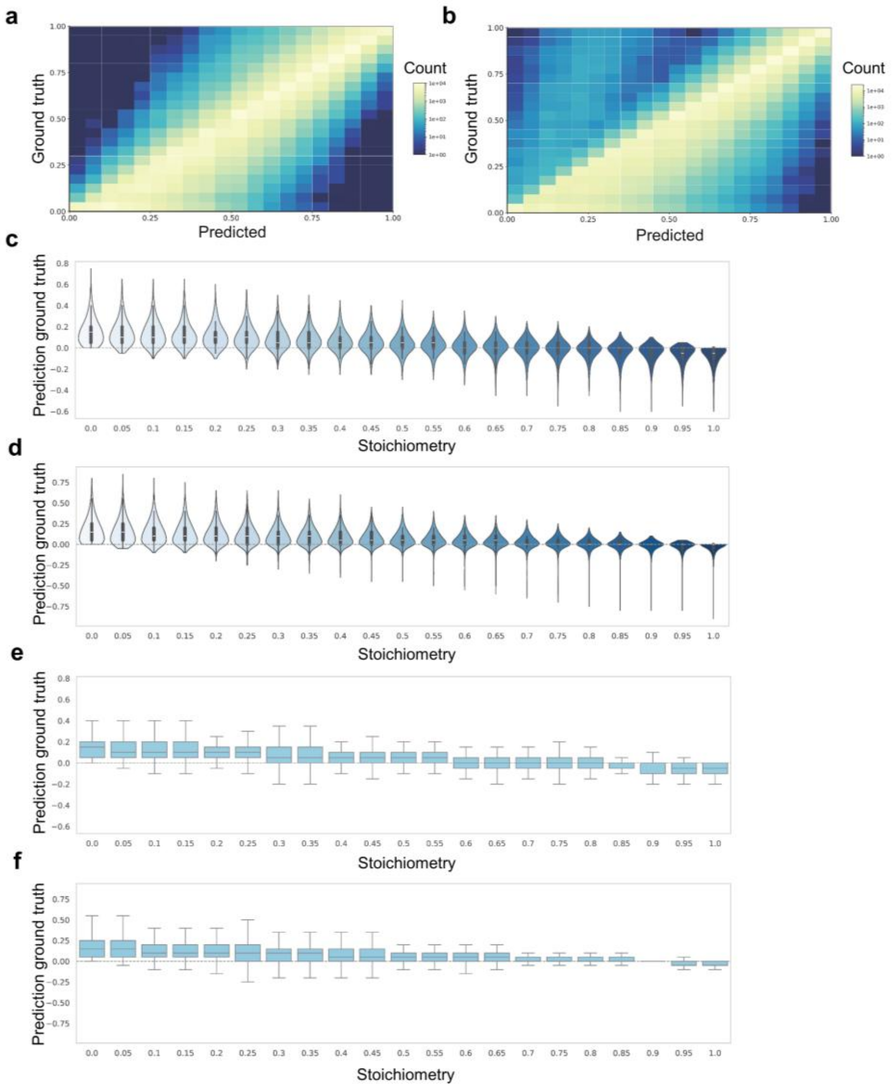


**Supplementary Fig.3 RedNano's capability for estimating modification rates. a** Correlation between predicted methylation stoichiometry by RedNano and the ground truth in the synthetic RNA dataset. **b** Correlation between predicted methylation stoichiometry by RedNano and the ground truth in the IVT dataset. **c** Deviation between predicted by RedNano and ground truth methylation stoichiometry in the synthetic RNA dataset. **d** Deviation between predicted by RedNano and ground truth methylation stoichiometry in the IVT dataset.

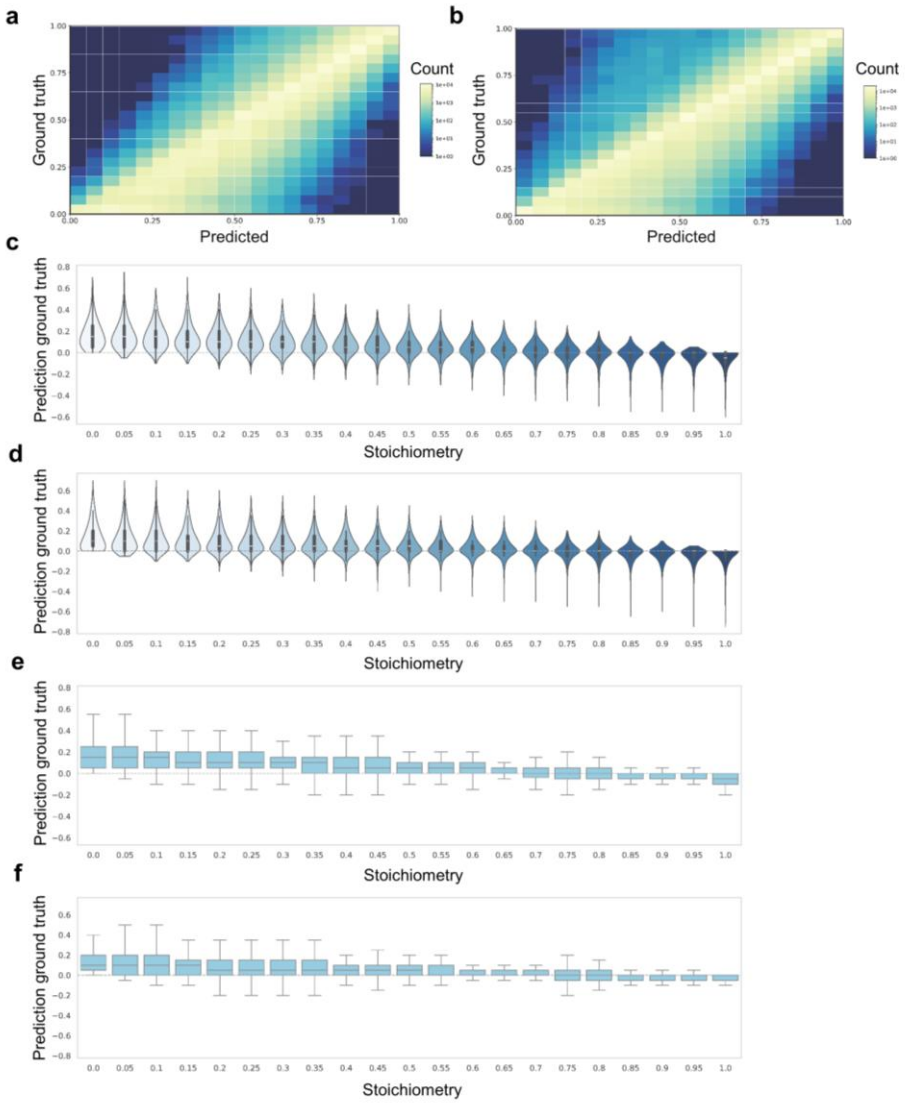


**Supplementary Fig.4 nanom6A's capability for estimating modification rates. a** Correlation between predicted methylation stoichiometry by nanom6A and the ground truth in the synthetic RNA dataset. **b** Correlation between predicted methylation stoichiometry by nanom6a and the ground truth in the IVT dataset. **c** Deviation between predicted by nanom6a and ground truth methylation stoichiometry in the synthetic RNA dataset. **d** Deviation between predicted by nanom6a and ground truth methylation stoichiometry in the IVT dataset.

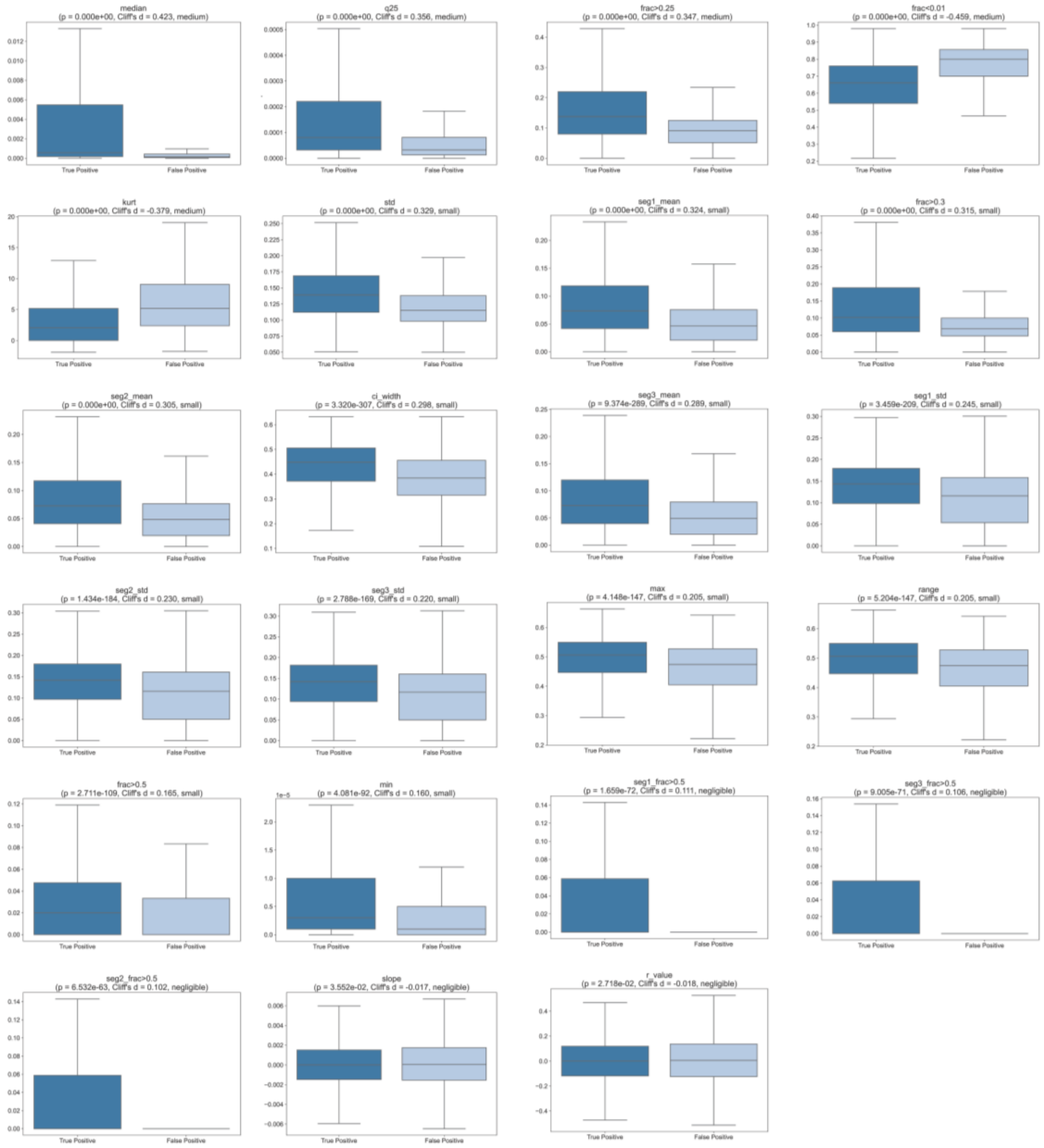


**Supplementary Fig.5 Statistical difference in read-level probability distributions between true and false positives where model trained on the synthetic RNA dataset.** Comparison of predicted read-level methylation distributions between true positives and false positives when the model is trained on the synthetic RNA dataset.

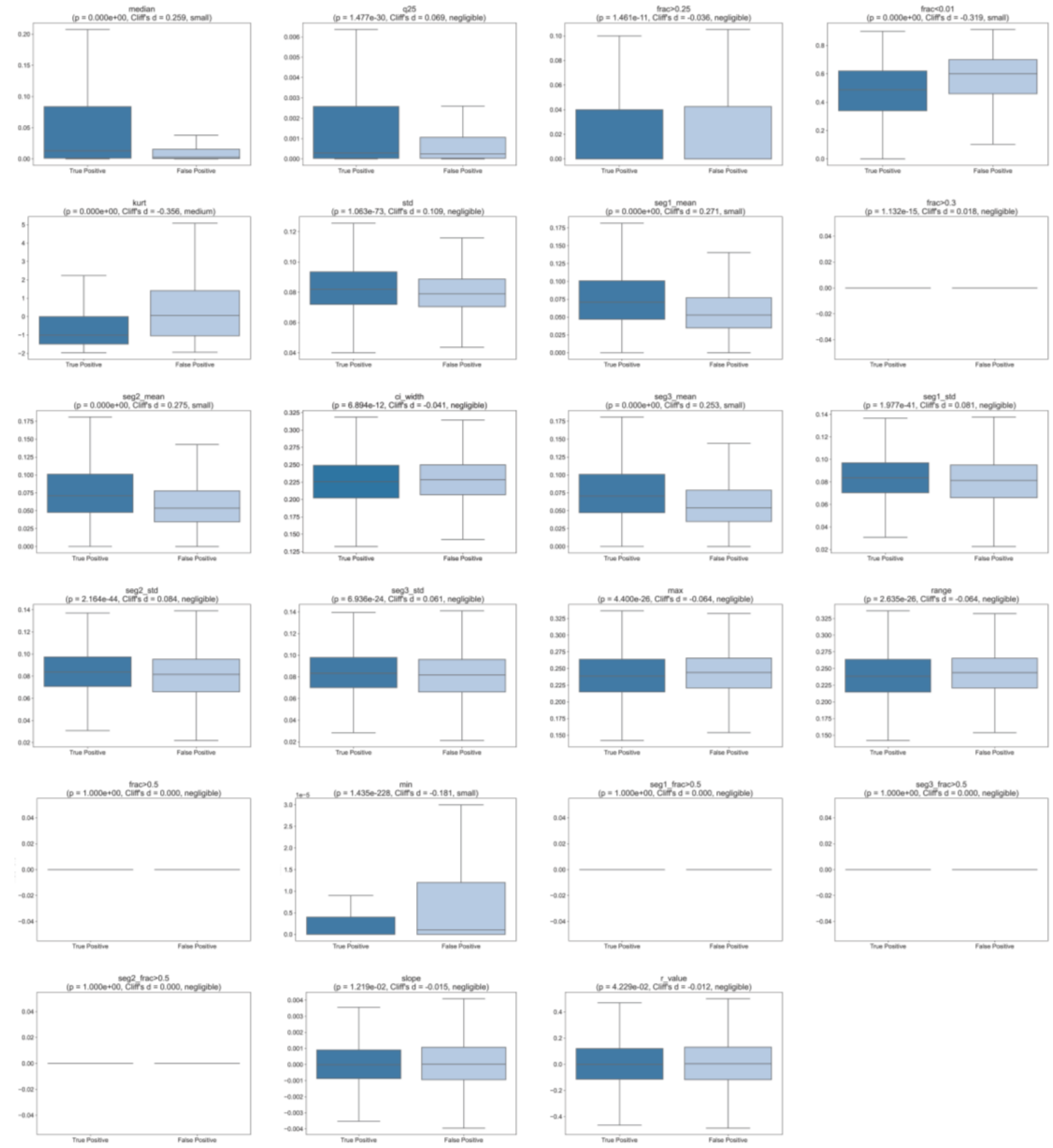


**Supplementary Fig.6 Statistical difference in read-level probability distributions between true and false positives where model trained on the *Arabidopsis* dataset.** Comparison of predicted read-level methylation distributions between true positives and false positives when the model is trained on the *Arabidopsis* dataset.

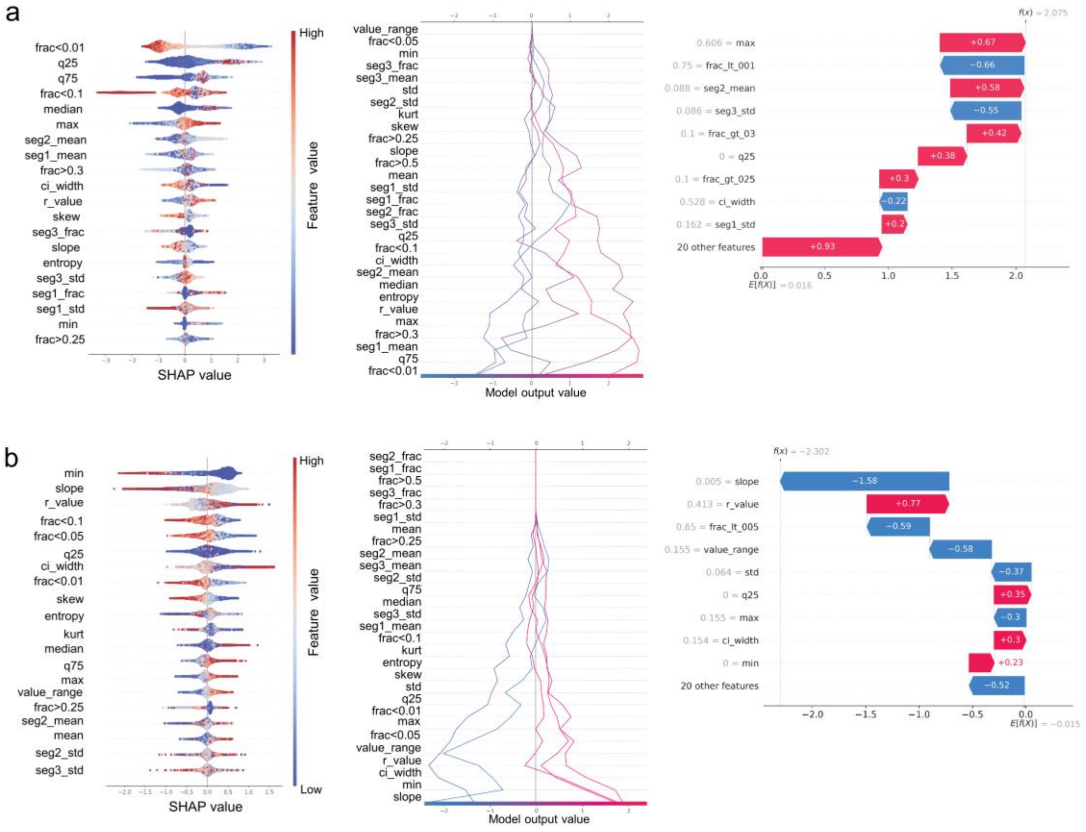


**Supplementary Fig.7 Model Explainability Analysis. a, b** SHAP-based interpretability of the model during cross-dataset transfer. The plots show feature contributions when transferring from the synthetic RNA dataset (a) and the *Arabidopsis* dataset (b) to the HEK293T dataset, respectively.

### Supplementary Tables

| Model | F1 | AUC | AUPR | Sensitivity | Specificity |
|---|---|---|---|---|---|
| MulitNano | 0.536 | 0.860 | 0.545 | 0.749 | 0.825 |
| RedNano | 0.517 | 0.853 | 0.530 | 0.772 | 0.797 |
| m6anet | 0.486 | 0.833 | 0.500 | 0.729 | 0.788 |
| nanom6a | 0.358 | 0.743 | 0.317 | 0.777 | 0.572 |
| EpiNano | 0.307 | 0.678 | 0.250 | 0.866 | 0.369 |

Supplementary Table S1. Performance of each model at the site-level on the HEK293T dataset in the DRACH motif.

| Model | F1 | AUC | AUPR | Sensitivity | Specificity |
|---|---|---|---|---|---|
| MulitNano | 0.544 | 0.859 | 0.558 | 0.783 | 0.786 |
| RedNano | 0.530 | 0.854 | 0.542 | 0.804 | 0.760 |
| m6anet | 0.497 | 0.829 | 0.512 | 0.757 | 0.747 |
| nanom6a | 0.370 | 0.734 | 0.340 | 0.816 | 0.492 |
| EpiNano | 0.330 | 0.660 | 0.261 | 0.891 | 0.316 |

Supplementary Table S2. Performance of each model at the site-level on the HEK293T dataset using the RRACH motif.

| Model | Pearson | MSE | MAE | $R^2$ |
|---|---|---|---|---|
| MulitNano | 0.983 | 0.00019 | 0.0099 | 0.935 |
| RedNano | 0.981 | 0.00021 | 0.0103 | 0.931 |
| m6anet | 0.975 | 0.00041 | 0.0146 | 0.862 |
| nanom6a | 0.924 | 0.00049 | 0.0152 | 0.837 |
| EpiNano | 0.734 | 0.00140 | 0.0275 | 0.536 |

Supplementary Table S3. Comparison between the predicted methylation results and ground truth for different motifs across models.

| Model | F1 | AUC | AUPR | Sensitivity | Specificity |
|---|---|---|---|---|---|
| MulitNano | 0.908 | 0.964 | 0.958 | 0.936 | 0.876 |
| RedNano | 0.892 | 0.955 | 0.945 | 0.945 | 0.828 |
| nanom6a | 0.890 | 0.951 | 0.940 | 0.930 | 0.842 |

Supplementary Table S4 Performance of each model at the read-level on the synthetic RNA dataset.

| Model | F1 | AUC | AUPR | Sensitivity | Specificity |
|---|---|---|---|---|---|
| MulitNano | 0.944 | 0.981 | 0.990 | 0.955 | 0.886 |
| RedNano | 0.927 | 0.967 | 0.979 | 0.954 | 0.829 |
| nanom6a | 0.919 | 0.960 | 0.973 | 0.957 | 0.793 |

Supplementary Table S5. Performance of each model at the read-level on the IVT dataset.

| Model | F1 | AUC | AUPR | Sensitivity | Specificity |
|---|---|---|---|---|---|
| MulitNano | 0.431 | 0.752 | 0.406 | 0.549 | 0.833 |
| RedNano | 0.423 | 0.737 | 0.391 | 0.473 | 0.873 |
| m6anet | 0.423 | 0.735 | 0.386 | 0.421 | 0.905 |
| nanom6a | 0.346 | 0.688 | 0.283 | 0.572 | 0.710 |
| EpiNano | 0.261 | 0.589 | 0.196 | 0.369 | 0.756 |

Supplementary Table S6. Results of transfer learning experiments: models trained on the synthetic RNA dataset and tested on the HEK293T dataset(DRACH motif).

| Model | F1 | AUC | AUPR | Sensitivity | Specificity |
|---|---|---|---|---|---|
| MulitNano | 0.551 | 0.833 | 0.508 | 0.624 | 0.875 |
| RedNano | 0.536 | 0.823 | 0.505 | 0.546 | 0.904 |
| m6anet | 0.470 | 0.818 | 0.495 | 0.388 | 0.948 |
| nanom6a | 0.340 | 0.682 | 0.305 | 0.720 | 0.508 |
| EpiNano | 0.249 | 0.600 | 0.239 | 0.230 | 0.866 |

Supplementary Table S7. Results of transfer learning experiments: models trained on the Arabidopsis dataset and tested on the HEK293T dataset(RRACH motif).

| Mestics | MultiNano | RedNano | nanom6A |
|---|---|---|---|
| Pearson | 0.951 | 0.932 | 0.930 |
| MSE | 0.010 | 0.015 | 0.016 |
| MAE | 0.071 | 0.087 | 0.090 |
| $R^2$ | 0.886 | 0.834 | 0.820 |

Supplementary Table S8. Model performance in methylation ratio prediction on the synthetic RNA dataset, evaluated by comparison with the ground truth modification ratios.

| Mestics | MultiNano | RedNano | nanom6A |
|---|---|---|---|
| Pearson | 0.957 | 0.938 | 0.929 |
| MSE | 0.008 | 0.013 | 0.017 |
| MAE | 0.053 | 0.074 | 0.083 |
| $R^2$ | 0.907 | 0.855 | 0.814 |

Supplementary Table S9. Model performance in methylation ratio prediction on the IVT dataset, evaluated by comparison with the ground truth modification ratios.

| Method | TP | FP | FN | TN | Precision | Recall |
|---|---|---|---|---|---|---|
| OurMethod | 29418 | 2914 | 24563 | 51074 | 0.910 | 0.545 |
| 0.6 | 31210 | 8334 | 22771 | 45654 | 0.789 | 0.578 |
| 0.7 | 24696 | 5354 | 29285 | 48634 | 0.822 | 0.458 |
| 0.75 | 21042 | 4059 | 32939 | 49929 | 0.838 | 0.390 |
| 0.8 | 17080 | 2899 | 36901 | 51089 | 0.855 | 0.316 |
| 0.9 | 7136 | 949 | 46845 | 53039 | 0.883 | 0.132 |

Supplementary Table S10. Performance of different models in false positives removal during transfer learning from the synthetic RNA dataset to the HEK293T dataset. Values represent results under different cutoff thresholds.

| Method | TP | FP | FN | TN | Precision | Recall |
|---|---|---|---|---|---|---|
| OurMethod | 21100 | 1735 | 32881 | 52253 | 0.924 | 0.391 |
| 0.5 | 26109 | 6614 | 27872 | 47374 | 0.798 | 0.484 |
| 0.6 | 22144 | 4218 | 31837 | 49770 | 0.840 | 0.410 |
| 0.7 | 18019 | 2638 | 35962 | 51350 | 0.872 | 0.334 |
| 0.75 | 15890 | 2024 | 38091 | 51964 | 0.887 | 0.294 |
| 0.8 | 13752 | 1528 | 40229 | 52460 | 0.900 | 0.255 |
| 0.9 | 8724 | 773 | 45257 | 53215 | 0.919 | 0.162 |

Supplementary Table S11. Performance of different models in false positives removal during transfer learning from the Arabidopsis dataset to the HEK293T dataset. Values represent results under different cutoff thresholds.
